# iCOMIC: a graphical interface-driven bioinformatics pipeline for analyzing cancer omics data

**DOI:** 10.1101/2021.09.18.460896

**Authors:** Anjana Anilkumar Sithara, Devi Priyanka Maripuri, Keerthika Moorthy, Sai Sruthi Amirtha Ganesh, Philge Philip, Shayantan Banerjee, Malvika Sudhakar, Karthik Raman

## Abstract

Despite the tremendous increase in omics data generated by modern sequencing technologies, their analysis can be tricky and often requires substantial expertise in bioinformatics. To address this concern, we have developed a user-friendly pipeline to analyze (cancer) genomic data that takes in raw sequencing data (FASTQ format) as input and outputs insightful statistics on the nature of the data. Our iCOMIC toolkit pipeline can analyze whole-genome and transcriptome data and is embedded in the popular Snakemake workflow management system. iCOMIC is characterized by a user-friendly GUI that offers several advantages, including executing analyses with minimal steps, eliminating the need for complex command-line arguments. The toolkit features many independent core workflows for both whole genomic and transcriptomic data analysis. Even though all the necessary, well-established tools are integrated into the pipeline to enable ‘out-of-the-box’ analysis, we provide the user with the means to replace modules or alter the pipeline as needed. Notably, we have integrated algorithms developed in-house for predicting driver and passenger mutations based on mutational context and tumor suppressor genes and oncogenes from somatic mutation data. We benchmarked our tool against Genome In A Bottle (GIAB) benchmark dataset (NA12878) and got the highest F1 score of 0.971 and 0.988 for indels and SNPs, respectively, using the BWA MEM - GATK HC DNA-Seq pipeline. Similarly, we achieved a correlation coefficient of r=0.85 using the HISAT2-StringTie-ballgown and STAR-StringTie-ballgown RNA-Seq pipelines on the human monocyte dataset (SRP082682). Overall, our tool enables easy analyses of omics datasets, with minimal steps, significantly ameliorating complex data analysis pipelines.

Availability: https://github.com/RamanLab/iCOMIC

## Introduction

Over the past couple of decades, genomic research has developed tremendously due to the rise of Next-Generation Sequencing (NGS) technologies. These rapid advances have had a considerable impact in the realm of sequence-based analysis: NGS has enabled researchers to discover novel DNA and RNA variants [1], along with differentially expressed genes [2]. Whole Genome/Exome Sequencing (DNA-Seq) pipelines identify nucleotide variants, while RNA Sequencing (RNA-Seq) enables quantification of gene expression. Whole Genome Sequencing further serves as a powerful tool in analyzing mutations in the context of cancer[3–4] and is also a bedrock for personalized medicine [5]. RNA-Seq further refines the essential interpretation of various biological phenomena.

Various bioinformatics tools have been developed to analyze the large amounts of data generated by NGS technologies. Data analysis poses a major hindrance to biologists, exacerbating the need for an automated pipeline. Even though new tools for genomic data analysis are being developed from time to time, a comprehensive toolkit does not exist. Extensive comparative studies that deal with different combinations of tools have been conducted [6] [7] [8]. Although software suites consisting of a combination of few tools exist [9–14], a user-friendly toolkit to aid non-programmers, incorporating a surplus number of bioinformatics tools, is missing [15]. Moreover, there is a dearth of open-source bioinformatics pipelines that enables comprehensive analysis of large (cancer) genomic datasets. [16].

To make life easier for clinical researchers and biologists, we developed iCOMIC, an open-source, standalone tool characterized by a Python-based Graphical User Interface and automated Bioinformatics pipelines for DNA-Seq and RNA-Seq data analysis. It serves as a point-and-click application facilitating genomic data analysis accessible to users with minimal programming expertise. iCOMIC provides a versatile, fully automated pipeline for the analysis of genomic data, with a user-defined combination of tools and a set of easily tunable parameters. In addition, iCOMIC grants users the privilege to customize pipelines in less than five simple steps, integrating a wide range of bioinformatics tools.

## Methods

The first step towards developing iCOMIC was to compile a set of tools following best practices for DNA-Seq and RNA-Seq analysis. We identified the most widely used tools for alignment of reads, variant calling, variant annotation, expression modeling, and differential expression analysis (Figure 1, Table 1, [6] [17]). Two of the most cited and widely used tools, FastQC [18] and Cutadapt [19], were chosen for quality control.

**Figure 1:**
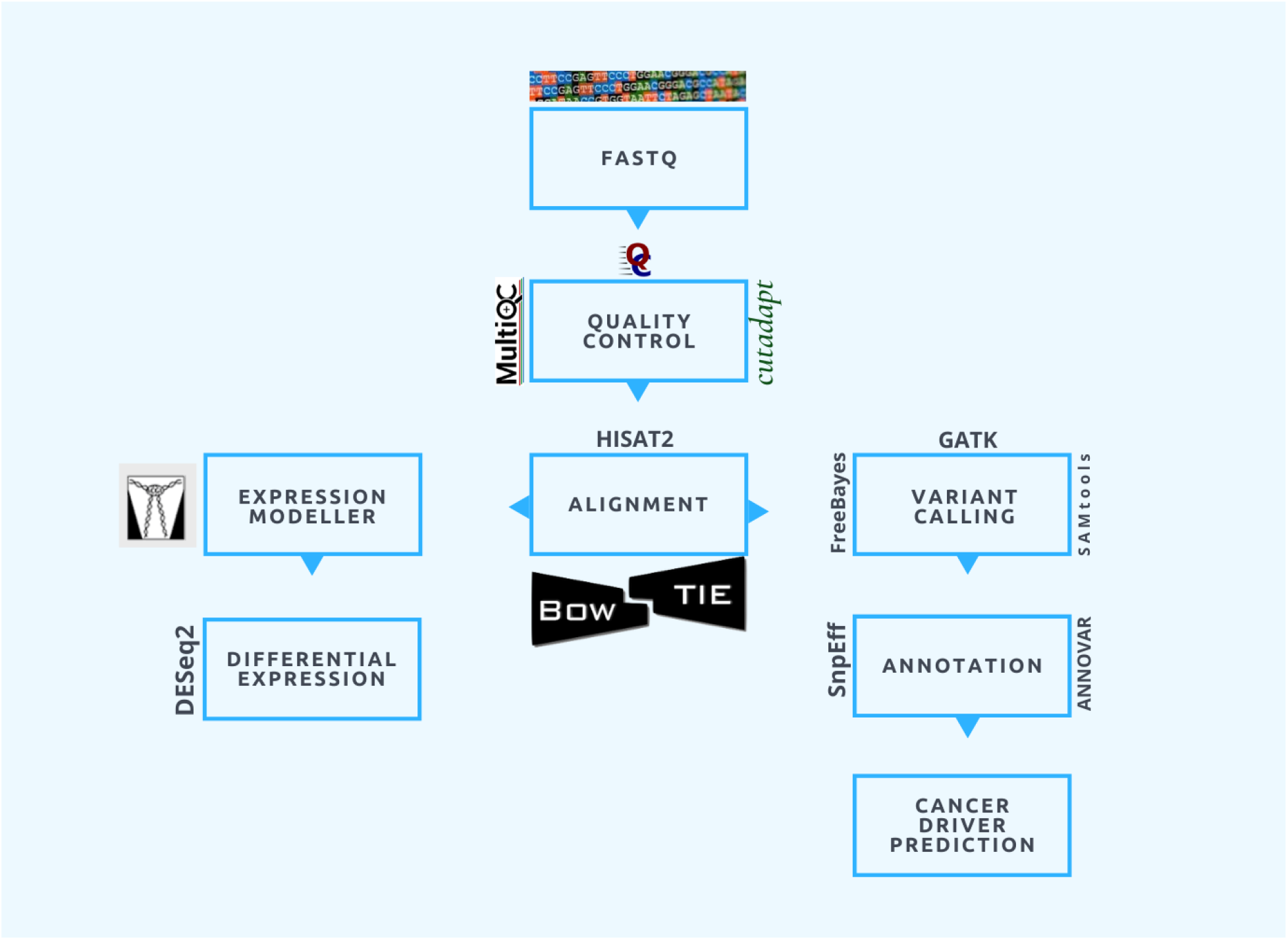
Schema for iCOMIC pipeline. Multiple workflows are embedded in iCOMIC providing users with the complete freedom to choose from the integrated tools. Both DNA-Seq and RNA-Seq pipelines take in raw FASTQ files as input. Quality control and alignment are common steps in both pipelines. FastQC and Cutadapt are the Quality control tools used and MultiQC is used to generate a consolidated report on Quality statistics. Analysis of RNA-Seq data includes mapping of sequencing reads to a reference genome using Aligner, Quantification of expression levels using Expression modeller and Differential expression analysis. On the other hand, steps in DNA-Seq analysis include Alignment followed by identifying the variants and annotating them. Tools incorporated in iCOMIC are listed in Table 1.

**Table 1.**
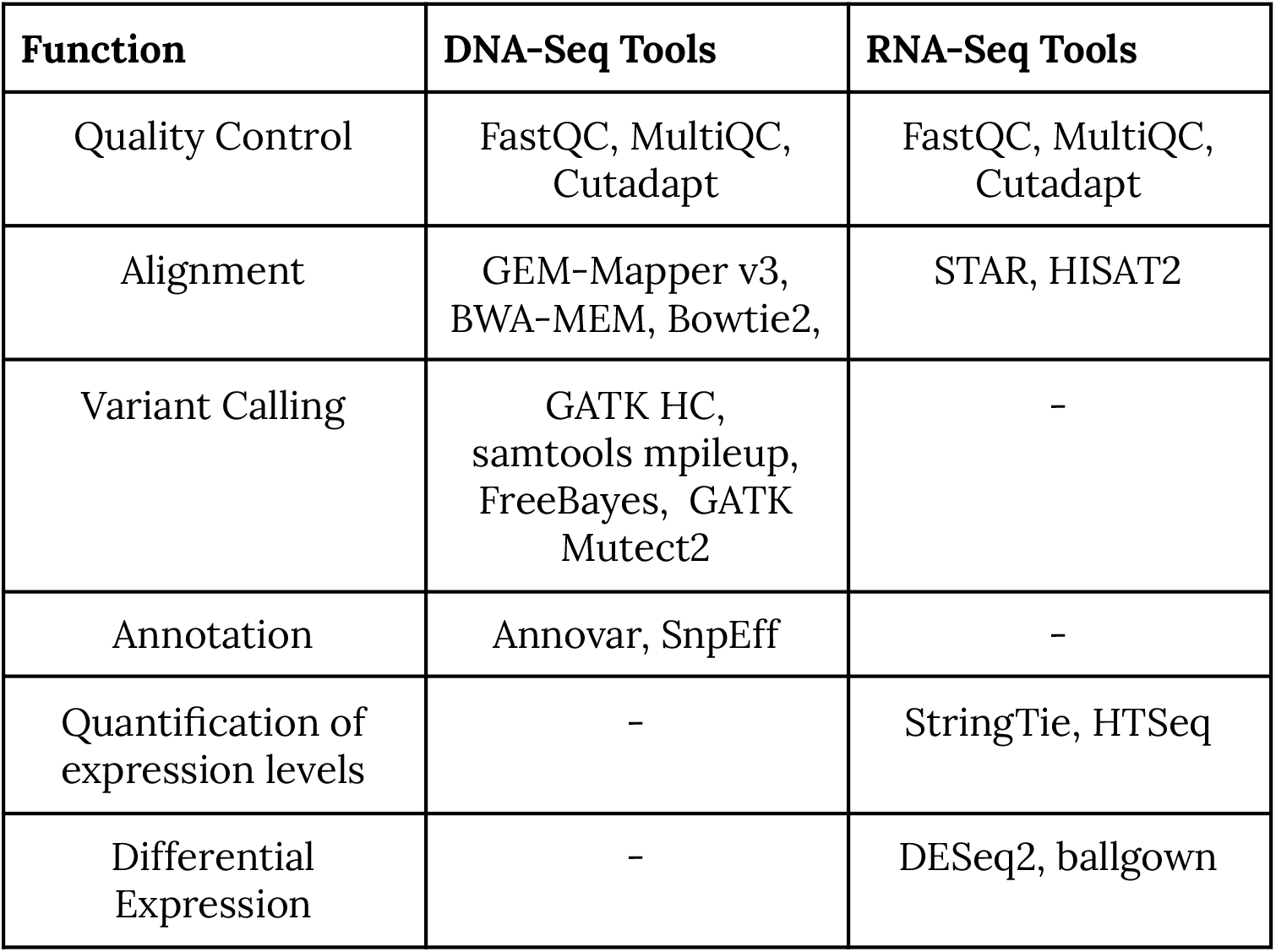
List of tools incorporated in iCOMIC along with their corresponding functions.

Aligner tools incorporated in iCOMIC for DNA-Seq data include BWA-MEM [20], GEM-Mapper v3 213], and Bowtie 2 [22]. Apart from the three major steps involved in analyzing whole-genome sequence data, several preprocessing steps are carried out intermediately following the GATK framework. This included sorting and indexing alignment files generated by the aligner, marking duplicates using Picard markduplicates[23], and base quality score recalibration followed by filtration of the variants called. Variant callers include GATK Mutect2 [24], samtools mpileup [25], FreeBayes [26], and GATK Haplotype-Caller [27]. Mutect2 identifies variants in normal-tumor pairs while the rest of the tools perform variant calling in a given sample analogous to the reference genome. SnpEff [28] and ANNOVAR [29] were the tools selected for variant annotation. The tool MultiQC [30] was used to aggregate the results obtained from the numerous tools in the workflow. In the case of whole genome/exome data, the MultiQC report was generated based on the results from tools such as FastQC and SnpEff. Two cancer-related data analysis tools, NBDriver [31], which uses a machine learning approach to identify the context of mutations, and cTaG [32], a tool to predict whether a given gene is a tumor suppressor (TSG) or an oncogene (OG), were also incorporated in the final pipeline(Figure 2).

**Figure 2:**
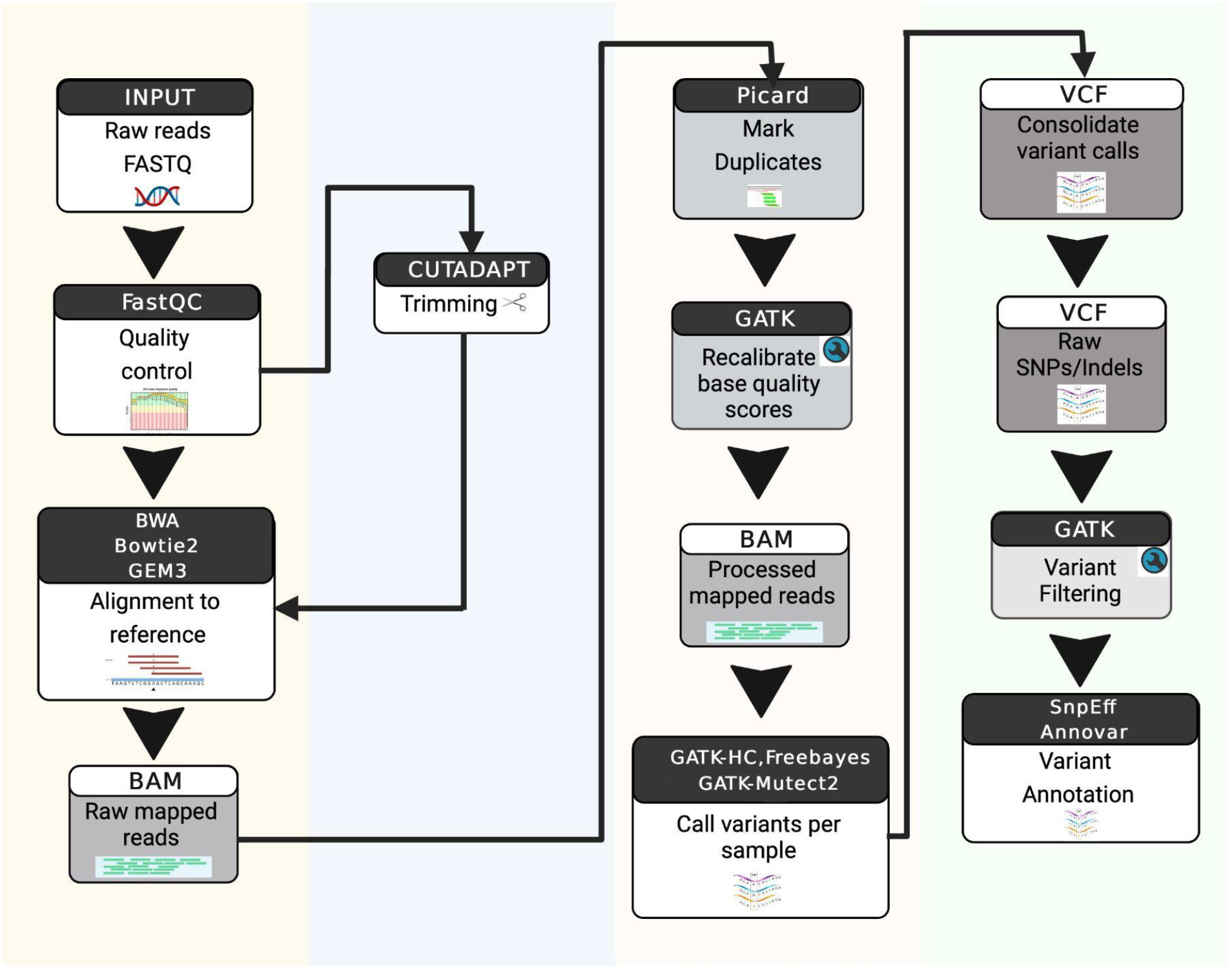
Schematic diagram of DNA-Seq pipeline. The input, followed by the application of various quality control techniques, alignment to the reference genome, variant calling, filtering and annotation are indicated in this figure.

For RNA-seq data, HISAT2 [33] and STAR [34] were selected to perform alignment following quality control. Expression modellers [6] such as StringTie [35] and HTSeq [36] were included to count the number of reads aligned to the genome. For differential expression analysis, ballgown [37] and DESeq2 [38] were incorporated. The MultiQC report for RNA-Seq data includes results from tools such as FastQC, Cutadapt, and STAR. Normalization is inherently carried out by the tool used for differential expression. (Figure 3).

**Figure 3:**
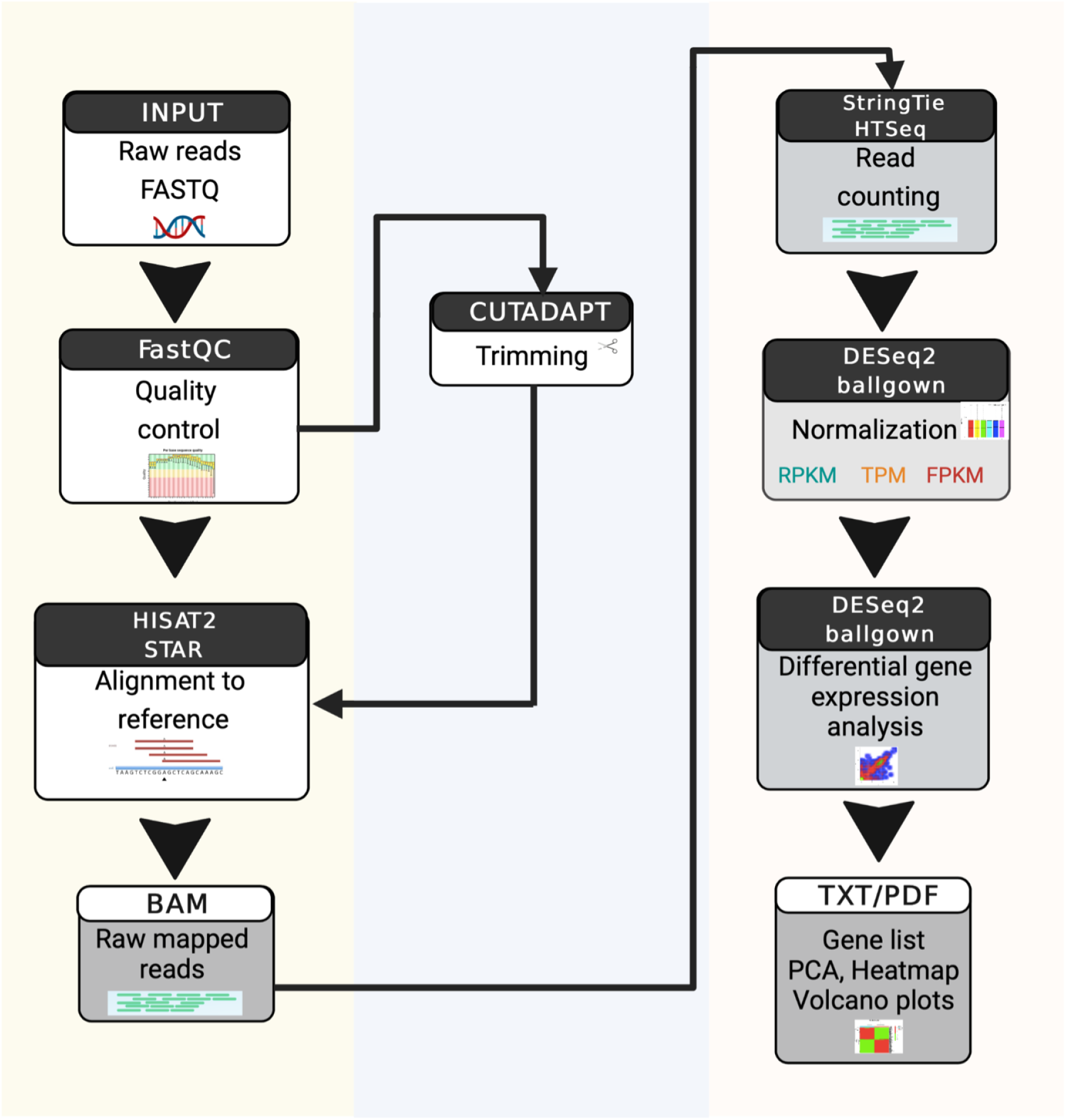
Schematic diagram of RNA-Seq pipeline. The input, followed by the application of various quality control techniques, alignment to the reference genome, counting the mapped reads, normalization, and differential expression analysis, ultimately generating the TXT/PDF output is detailed in this figure.

iCOMIC is embedded in the popular workflow management system, Snakemake [39]. The analysis workflow has ‘rules’ as the building blocks, which describe the connection between the input and output [40] (Figure 4). It enables easy connectivity between the different tools/software within the workflow. The ‘rules’ specify input files, output files, log files, and wrapper/shell commands. The Snakemake wrapper scripts together with the conda environment manage the automated installation of software and their dependencies. Tools without a wrapper script are configured separately, and shell commands are used for its execution. According to the user’s selection, appropriate rules are combined in a ‘Snakefile’ to generate the target output. The input file paths and parameters set by the user are automatically fed into a configuration file, referred to as the ‘config file.’ Multiple config files and Snakefiles are auto-generated for the quality check of the input data, generating genome index, and executing the main workflow.

**Figure 4:**
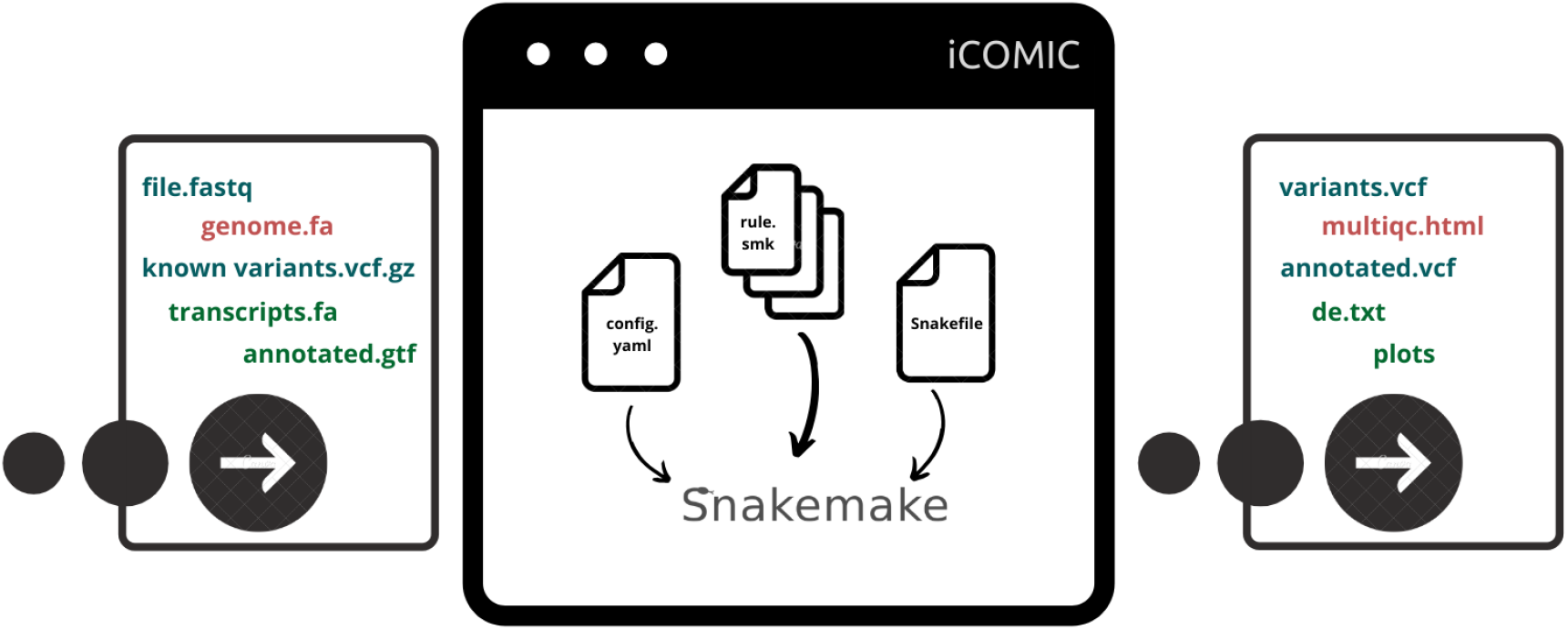
Snakemake workflow management system. All the input and output files in blue colour are those corresponding to DNA-Seq analysis and those in green correspond to RNA-seq analysis. The common files for DNA and RNA-Seq analysis are represented in red. ‘Rule’ files specifying the input, output and the shell/wrapper script form the basic units of Snakemake. Each rule corresponds to individual tools. The additional parameters for the tools are indicated in the ‘config’ file. According to the choice of tools made by the user, rules are integrated into the Snakefile and the workflow is executed.

We implemented the iCOMIC pipeline in the form of a Graphical User Interface (GUI). The GUI has been developed using PyQt5, a Python binding of the cross-platform GUI toolkit Qt. The GUI retrieves the user input files and the parameters and communicates with the Snakemake rules to set up the analysis. A Python wrapper binds together the PyQt5 GUI and the Snakemake workflow of iCOMIC. After completing the initial requirements, the iCOMIC GUI can be accessed using a single command, ‘`icomic’.`

## Results and Discussion

### Major features of the pipeline

NGS data has become an indispensable tool for biological research, although the data analysis can be daunting for non-bioinformaticians. iCOMIC has been introduced to overcome this concern to a certain extent. It serves as a stand-alone end-to-end analysis toolkit for DNA-Seq and RNA-Seq data. The DNA-Seq component of iCOMIC supports both germline and somatic variant calling. In conventional analysis pipelines like Galaxy, workflows need to be built, whereas iCOMIC has various inbuilt pipelines that automatically transfer output from one tool to the next. iCOMIC provides an interactive and user-friendly GUI, specifically created to accommodate users with minimal programming expertise. On another note, iCOMIC allows expert bioinformaticians to perform analysis incorporating additional tools and advanced parameters, saving time building the pipeline. The steps to be followed for writing new rules for integrating additional tools are detailed in the documentation at https://icomic-doc.readthedocs.io/en/latest/. The GUI is embedded in a Python wrapper script that connects the Snakemake workflows. Users can select an array of tools from the predesigned combinations best suited for their requirements. The best connectivity between the tools has been taken into account to design these individual workflows. iCOMIC provides the user with the necessary means to replace modules or alter the pipeline. Furthermore, the conda environment ensures easy installation of tools and dependencies.

### Prediction of tumour suppressor genes and oncogenes using the cTaG algorithm

The cTaG (classify TSG and OG) model [32] identifies driver genes by classifying them as tumor suppressor genes (TSGs) and oncogenes (OGs). Given a cohort of samples, the pan-cancer model calculates ratio-metric features from somatic mutation data, capturing mutations’ functional impact. Unlike other computational methods that use background mutation rate (BMR) to identify genes with a higher mutation rate as driver genes, cTaG captures the effect of a mutation on the gene’s functionality. Methods using BMR are biased towards genes with high mutation rates [41], and we know that while few driver genes have a high mutation rate, most do not. The mutations in TSG and OG differ; we found nonsense mutations more commonly found in TSGs than OGs. The cTaG method contains binary classifiers that classify genes as TSG or OG. The genes are labeled as TSGs or OGs based on consensus across various models.

To build a pan-cancer model, the model was trained on somatic mutation data from COSMIC (v79) [42] from different cancer types. We used ratio-metric and entropy features to classify genes as TSG or OG. The cTaG model uses the random forest classification algorithm to generate the pan-cancer model. A filtered list of genes from the Cancer Gene Census (CGC; [42, 43]) is used to label genes as TSG or OG, used to train the pan-cancer model. The pan-cancer model successfully identified tissue-specific driver genes when employed on a cohort from single tissue of origin. The method takes a single `maf` file with annotated mutations for the cohort of samples as input and generates each gene’s ratio-metric and entropy features. Additional arguments such as the percentile and threshold to define highly-mutated samples can be specified. The percentile argument defines the top percentile genes to be considered from predictions made by each model for the final consensus. The default value is 5. Highly mutated samples are skipped from the analysis. By default, samples with more than 2000 mutations are omitted during analysis. The cTaG model labels these genes as TSG and OG based on the consensus across the random forest models. The model returns the list of all genes and their labels predicted by each model, along with its presence in the top percentile. Our method identifies genes with high as well as low mutation rates. The pan-cancer model also predicts tissue-specific driver genes.

### Prediction of driver and passenger mutations using NBDriver

Differentiating between driver and passenger mutations from sequenced cancer genomes is essential to targeted therapy and precision medicine. Despite the dramatic advances in developing predictive algorithms to differentiate between driver and passenger mutations, very few have concentrated on utilizing the local sequence context as potential features for further analysis. To capture this information, we built a robust machine learning model called NBDriver [31], which uses raw nucleotide sequences surrounding cancer-causing mutations as features to build machine learning models.

Our training data consisted of missense mutations from 58 genes containing experimentally validated functional impacts from several studies. To obtain a numerical representation of the sequence features surrounding the mutations, we used commonly used natural language processing tools such as the TF-IDF Vectorizer, the Count Vectorizer, and the One-Hot Encoder. Using kernel density estimation techniques, we showed that the underlying distributions of the neighbourhood sequences surrounding driver and passenger mutations are significantly different from one another. We utilized this information to build robust machine learning models using a repeated cross-validation strategy and report the median values of the performance metrics for each feature-classifier pair. To increase the prediction performance, we integrated sequence features derived from raw nucleotide sequences with other genomic, structural, and evolutionary features, resulting in the development of a pan-cancer mutation effect prediction tool, NBDriver, which was highly efficient in identifying pathogenic variants from five independent validation datasets. An ensemble predictor containing NBDriver, CONDEL [44], and MutationTaster [45] outperformed existing pan-cancer models in prioritizing a literature-curated list of driver and passenger mutations. Considering only the true positive mutation predictions from NBDriver, we identified a list of 138 driver genes with known functional evidence from multiple sources. Overall, our study underpinned the efficacy of utilizing raw nucleotide sequences as features for building robust machine learning models to distinguish between driver and passenger mutations.

### Architecture of the GUI

The iCOMIC GUI has four major tabs. The first tab is for whole-genome/exome sequence analysis to identify annotated variants, and the second is for RNA-Seq analysis to identify differentially expressed genes. Each tab has a set of sub-tabs that allows the users to input necessary information from their end.

The sub-tabs include:

1. Input Data: Enables the user to provide raw FASTQ files, genome fasta files, and known variants in the Whole Genome/Exome Sequencing pipeline. Additionally, for the RNA-Seq pipeline, the ‘Input Data’ tab contains fields to input FASTQ files, genome fasta files, and annotated files to enable further analysis. The user can navigate through the tabs with the ‘Next’ and ‘Previous’ buttons. The “Logs” tab at the bottom of each tab in the GUI displays the commands executed by the user. This checks the input data for any inconsistencies and then proceeds with the analysis.
2. Quality Control: The Quality Control tab allows the user to perform sequence quality analysis with FastQC, following which reads can be trimmed if needed using Cutadapt for both DNA-Seq and RNA-Seq pipelines. A MultiQC report aggregating the results of the FastQC output will be displayed. The “Progress Bar” allows the user to visualize the progress of the analysis.
3. Tool Selection: This allows the user to select aligners, variant callers, and annotation tools for DNA-Seq data and aligners, expression modelers, and differential expression analysis tools for RNA-Seq data. Users can input parameters, provide a genome index for aligners, and other necessary inputs for each tool. The user is free to upload or generate the genome index files to shorten computation time. Index for the *Homo sapiens* genome build GRCh37 (hg19) and GRCh38 (hg18) corresponding to each given aligner is provided together with the iCOMIC package.
4. Run: Permits the user to perform analysis based on the input information. Depending on the selection of tools, the particular pipeline will run.Results: This tab enables users to visualize output statistics of the tools used, and the major output files, specifically, the variants called and the annotated variants in the case of DNA-Seq pipeline, and differentially expressed genes for RNA-Seq pipeline.

The third tab is for running the cTaG tool, which is used to identify tumor suppressor genes (TSGs) and oncogenes (OGs). The user needs a `maf` file to run the tool along with the necessary parameters. The user can also generate a `maf` file from the `vcf` file of the DNA-Seq result. The fourth tab is for running the NBDriver tool used to differentiate between driver and passenger mutations; this again takes as input a `vcf` file.

### Benchmarking of tools

#### Evaluating the performance of germline variant calling pipelines

Performance validation of the germline variant calling workflows available with iCOMIC was done using Genome In A Bottle (GIAB) benchmark sets, based on Zook *et al.* [46]. To this end, we ran 18 different combinations of aligners, germline variant callers, and annotators integrated with iCOMIC on the widely used benchmark dataset NA12878/HG001 to obtain separate `vcfs` (Table 2). Generated `vcfs` were then compared with GIAB/NIST HG001 v2.19 truth data [47], restricting the comparison to the GIAB v2.19 BED file coordinates. We adopted the benchmark framework developed by the Global Alliance for Genomics and Health (GA4GH) benchmarking team [48] for `vcf` comparison. The method involved the generation of an intermediate `vcf` with a standardized variant representation using `Vcfeval` by Real-Time Genomics (RTG) tools [49] followed by quantification of performance metrics using `qfy.py`, which is a part of `hap.py` benchmarking toolkit [50]. The summary of accuracy measures is highlighted in Table 2. The analysis was performed, specifying ten threads for running each tool for all the workflow combinations. iCOMIC provided the highest F1 score of 0.971 and 0.988 for indels and SNPs, respectively, for the BWA MEM - GATK HC pipeline among all the combinations (Table 2).

**Table 2:** Summary of Germline variant benchmarking with NA12878/HG001 dataset.

| Workflow | Type | Recall | Precision | F1 score |
| --- | --- | --- | --- | --- |
| BWA MEM-GATK HC-SnpEff | INDEL | 0.967 | 0.976 | 0.971 |
|  | SNP | 0.978 | 0.998 | 0.988 |
| BWA MEM-freebayes-SnpEff | INDEL | 0.931 | 0.917 | 0.924 |
|  | SNP | 0.979 | 0.997 | 0.988 |
| BWA MEM-GATK HC-Annovar | INDEL | 0.967 | 0.976 | 0.971 |
|  | SNP | 0.978 | 0.998 | 0.988 |
| BWA MEM-Bcftools-Annovar | INDEL | 0.741 | 0.838 | 0.789 |
|  | SNP | 0.976 | 0.996 | 0.986 |
| Gem3-GATK HC-SnpEff | INDEL | 0.964 | 0.978 | 0.971 |
|  | SNP | 0.977 | 0.999 | 0.988 |
| Gem3-Freebayes-SnpEff | INDEL | 0.934 | 0.92 | 0.927 |
|  | SNP | 0.978 | 0.998 | 0.988 |
| BWA MEM-GATK<br>HC-Annovar | INDEL | 0.967 | 0.976 | 0.971 |
|  | SNP | 0.978 | 0.998 | 0.988 |
| BWA<br>MEM-Freebayes-Annovar | INDEL | 0.931 | 0.917 | 0.924 |
|  | SNP | 0.979 | 0.997 | 0.988 |
| Gem3-GATK HC-Annovar | INDEL | 0.964 | 0.978 | 0.971 |
|  | SNP | 0.977 | 0.999 | 0.988 |
| Gem3-Freebayes-Annovar | INDEL | 0.934 | 0.92 | 0.927 |
|  | SNP | 0.978 | 0.998 | 0.988 |
| BWA MEM-Bcftools-SnpEff | INDEL | 0.741 | 0.838 | 0.789 |
|  | SNP | 0.976 | 0.996 | 0.986 |
| Gem3-Bcftools-SnpEff | INDEL | 0.781 | 0.353 | 0.486 |
|  | SNP | 0.975 | 0.997 | 0.986 |
| Bowtie2-GATK HC-SnpEff | INDEL | 0.847 | 0.978 | 0.908 |
|  | SNP | 0.953 | 0.998 | 0.975 |
| Bowtie2-GATK HC -Annovar | INDEL | 0.847 | 0.978 | 0.908 |
|  | SNP | 0.953 | 0.998 | 0.975 |
| Bowtie2-Freebayes-SnpEff | INDEL | 0.717 | 0.909 | 0.802 |
|  | SNP | 0.945 | 0.996 | 0.97 |
| Bowtie2-Freebayes -Annovar | INDEL | 0.717 | 0.908 | 0.802 |
|  | SNP | 0.945 | 0.996 | 0.97 |
| Bowtie2-Bcftools-SnpEff | INDEL | 0.648 | 0.891 | 0.75 |
|  | SNP | 0.944 | 0.985 | 0.964 |
| Bowtie2-Bcftools-Annovar | INDEL | 0.648 | 0.891 | 0.75 |
|  | SNP | 0.944 | 0.985 | 0.964 |

#### Evaluating the performance of RNA-Seq pipelines

The validation of the performance of RNA-Seq workflows available with iCOMIC was done using the Human monocyte RNA-Seq dataset from NCBI-SRA (SRP082682) [6]. We ran four different combinations of aligners, expression modellers, and differential expression tools integrated with iCOMIC on the RNA-Seq benchmark dataset to obtain differentially expressed genes between classical and non-classical monocytes. Comparison of differentially expressed genes between the genes identified and the non-classical and classical monocytes from the reference microarray dataset (GSE34515) obtained from the NCBI GEO database was performed. The fold change correlation [51] was calculated between reference microarray and RNA-Seq data. Based on the fold change correlation computed for the four different pipelines, it is evident that HISAT2-StringTie-ballgown and STAR-StringTie-ballgown pipelines performed the best with a correlation coefficient of r=0.85, higher than the values obtained for HISAT2-HTSeq-DESeq2 and STAR-HTSeq-DESeq2 pipelines (Figure 5).

**Figure 5:**
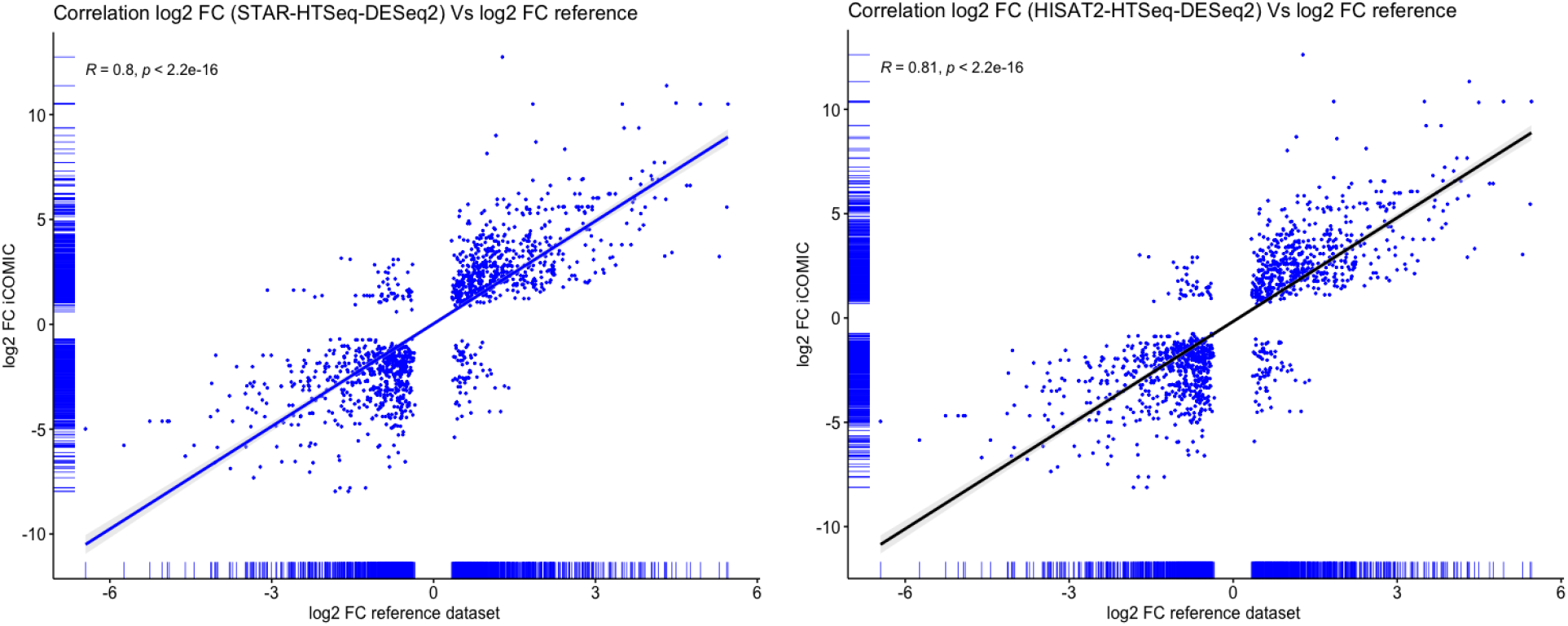
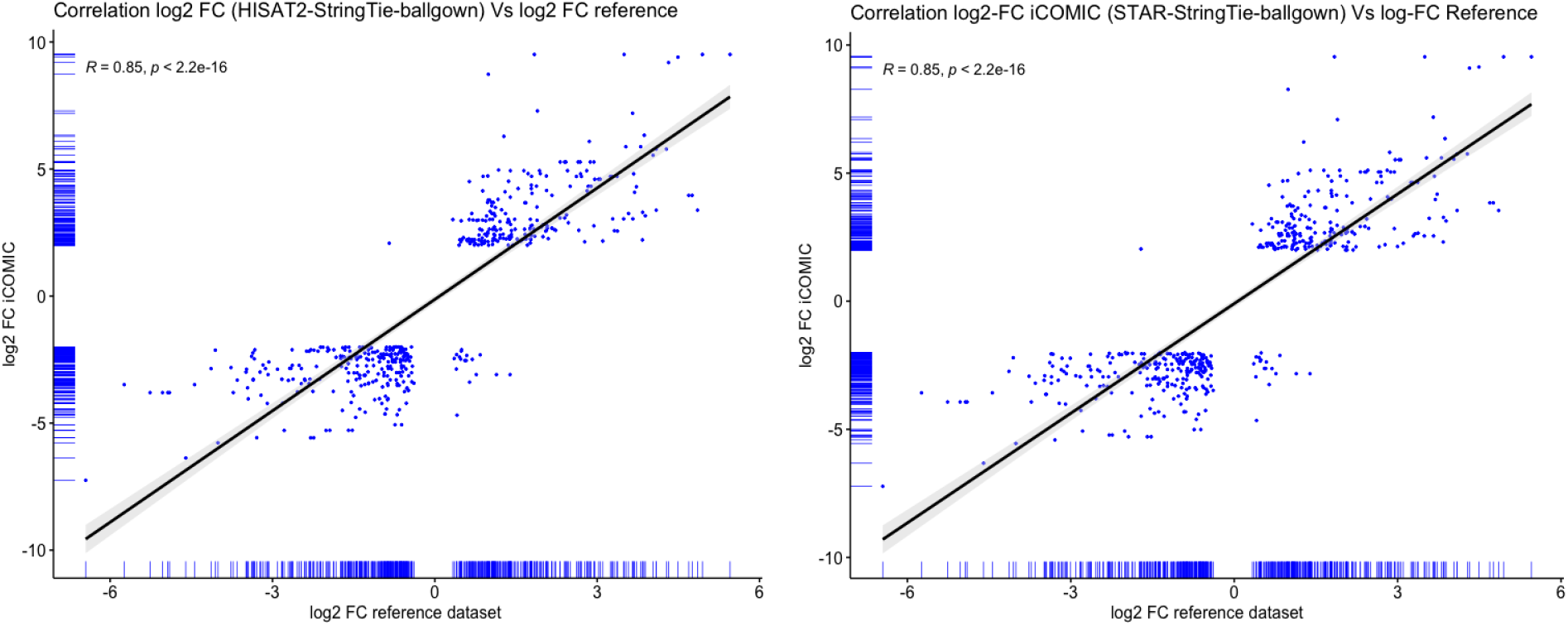
Fold change correlation between iCOMIC and reference dataset for the four workflows. The Pearson correlation coefficient was used to calculate fold changes.

#### Performance comparisons between iCOMIC and GALAXY

Galaxy [52] is a popular publicly available web-based interface consisting of many tool combinations and workflows to perform various studies. It offers ease of access to complex computational analyses to users with minimal programming expertise and the functionality to examine large datasets in a multi-step process. Considering the similarities in features between iCOMIC and Galaxy, pipeline validation was conducted using the same methods to establish a comparison scale for performance.

Pipelines are fully automated in iCOMIC and the most predominantly used workflows are pre-built. While workflows need to be specified in Galaxy in order to automate, iCOMIC comes with a set of preinstalled pipelines for both DNA and RNA-Seq analysis. In iCOMIC, the user has the option to specify a large number of files in the form of a table, while no such provision is available on Galaxy. Although extensive resources and tutorials are provided for the use of Galaxy, we believe it has a steeper learning curve than iCOMIC, which is mostly a point-and-click type application. Certain tools are unavailable on the Galaxy server and need to be installed from the tool shed, while iCOMIC installations are readily done via the conda environment, with minimal user input/interference.

While analysing multiple samples on a workflow in Galaxy, it is required to rename the files prior to passing it as input to the next tool in the pipeline which can be tedious when there is a large number of samples. On the other hand with iCOMIC, the entire process is automated. Undoubtedly, Galaxy has its advantages, and is a very popular pipeline for genomic data analysis. Yet, iCOMIC excels in its simplicity, and we believe it will be more inviting for biologists and clinical researchers to quickly analyse their data.

##### Comparison in terms of germline variant calling

DNA-Seq analysis was performed using BWA-MEM, Freebayes, and SnpEff for each step in the analysis workflow using Genome In A Bottle (GIAB) dataset. Performance validation for the same was done using a process similar to that of iCOMIC. The variant files obtained as output from this pipeline were compared with GIAB/NIST HG001 v2.19 truth data restricting the comparison to the GIAB v2.19 BED file coordinates. `Vcfeval` method was used for vcf comparison, and the quantification of performance metrics was computed using the `hap.py` algorithm. Comparison of the performance metrics such as F1 and precision scores between iCOMIC and Galaxy indicates that the values are pretty similar. Considering the SNP F1 score, iCOMIC has 0.988 and Galaxy has 0.980, proving that the performance of iCOMIC is on par with that of Galaxy (Table 3).

**Table 3:** Summary of Germline variant benchmarking with NA12878/HG001 dataset using Galaxy.

| Workflow | Type | Recall | Precision | F1 score |
| --- | --- | --- | --- | --- |
| BWA MEM-freebayes-SnpEff | INDEL | 0.887 | 0.948 | 0.917 |
|  | SNP | 0.976 | 0.984 | 0.980 |

##### Comparison in terms of differential expression analysis

RNA-Seq analysis was performed using the tools STAR, HTSeq, and DESeq2 for each step in the analysis workflow using the Human monocyte RNA-Seq dataset from NCBI-SRA (SRP082682). The fold change correlation was calculated between the reference microarray dataset (GSE34515) obtained from the NCBI GEO database and RNA-Seq data for iCOMIC and Galaxy. The values were found to be 0.8 and 0.66, respectively. Comparison of the fold change correlation between iCOMIC and Galaxy indicates that the value of iCOMIC is higher than that of GALAXY in the particular pipeline (Figure 6).

**Figure 6:**
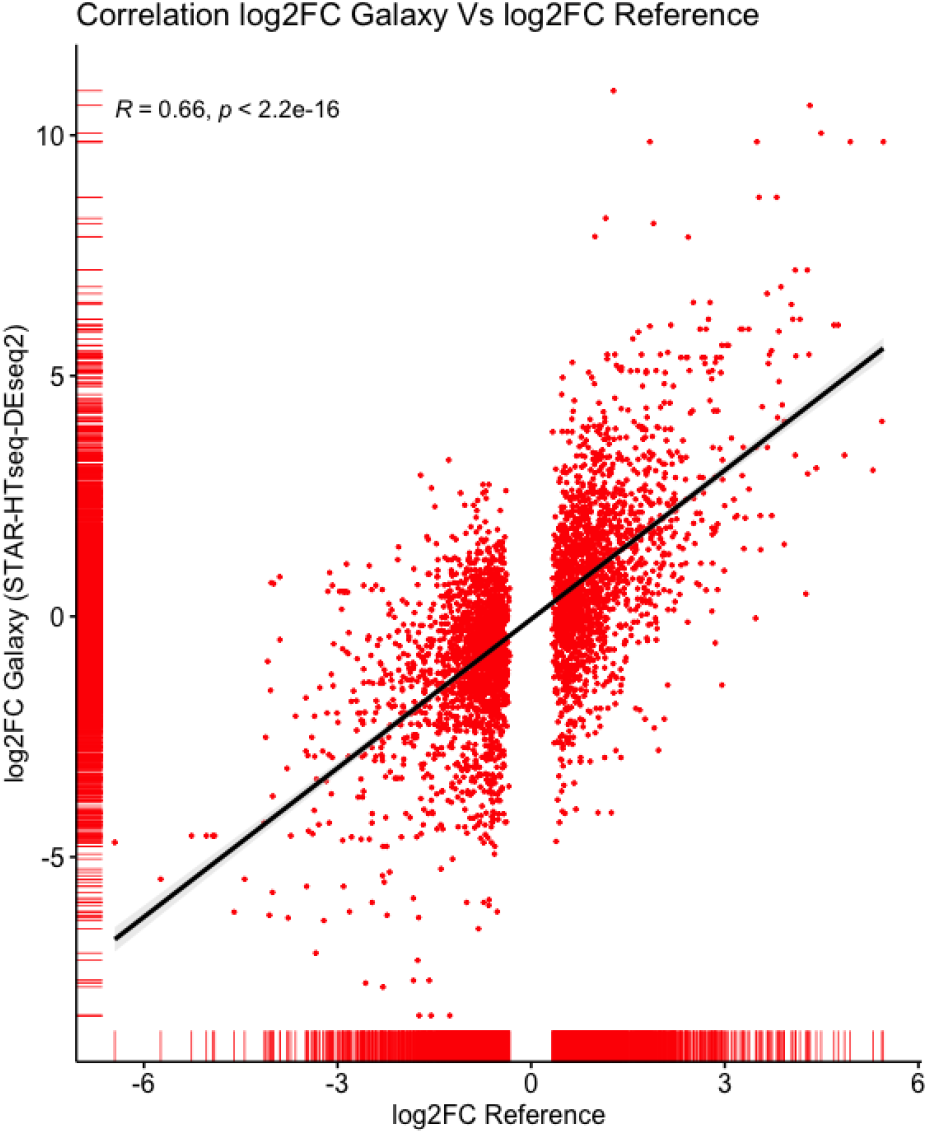
Fold change correlation between Galaxy and reference dataset for STAR-HTSeq-DESeq2 workflow. The Pearson correlation coefficient was used to calculate fold changes.

### Feature comparison between iCOMIC and other bioinformatics pipelines

There exist many pipelines that integrate different bioinformatics tools for genomic data analysis. A comprehensive feature comparison with previously developed workflows (snakePipes [8], Sequanix [40], Omics Pipe [7], GenPipes [53], CANEapp [15], ARMOR [54], Galaxy [52], VIPER [55], systemPipeR [56], CLC Genomics Workbench [16]) was performed to highlight the significance of iCOMIC (Table 4). The accessibility of the pipeline through GUI, availability to the public, cloud support, the ability of automated execution of an entire pipeline, user freedom to choose tools of interest, and programming skills required for the analysis include some of the features used for comparison. Only those tools that share common features with iCOMIC were considered for comparison. In addition, a comparison with the most popular bioinformatics tools like Galaxy [52] and CLC genomics workbench was performed. Galaxy, one of the customary scientific platforms for bioinformatics data analysis, does not afford inbuilt pipelines, whereas CLC genomics workbench is not open source. The comparative analysis highlights the open-source GUI with end-to-end automated analysis integrating a plethora of tools as the strongest aspects of iCOMIC.

**Table 4:**
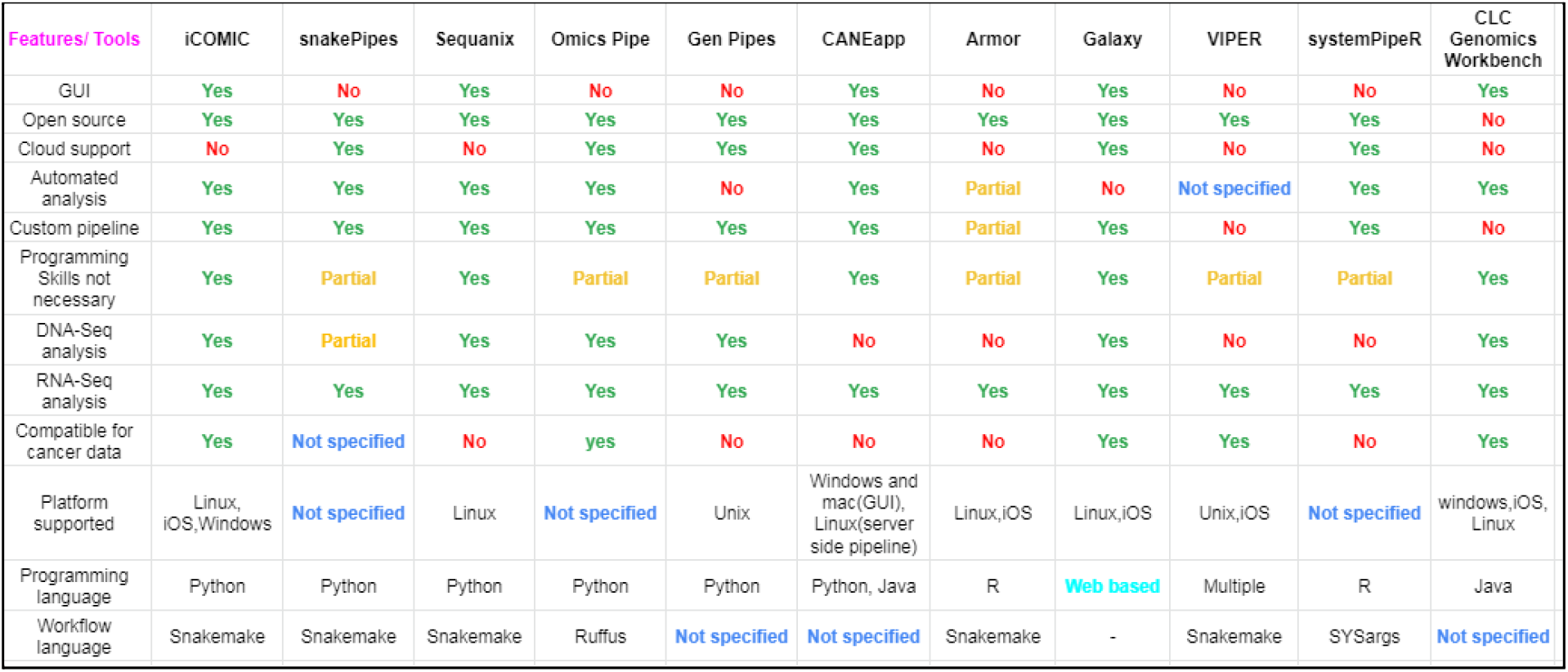
Comparison of iCOMIC with existing tools for genomic data analysis. ‘Yes’ specifies the presence of the feature, ‘No’ indicates the absence of a feature, ‘Partial’ indicates the presence of some aspects of the particular feature, and ‘Not Specified’ indicates that the information is not available. The features which are compared included: 1) The accessibility of the tool through Graphical User Interface, 2) commercial availability of the tool, 3) ability of the tool to run on the cloud, 4) automated analyses of the pipeline, 5) ability of the user to customize their own pipeline, 6) programming skills required for performing the analysis, 7) availability of DNA-Seq analysis pipeline, 8) availability of RNA-Seq analysis pipeline, 9) compatibility of the tool for cancer data. Furthermore, we have also listed out the platforms supported by the tools, programming language used to build the tool and workflow language used to write pipelines.

## Conclusion

Here, we present iCOMIC to analyze genomic data quickly. Analyzing the large amount of data generated by Next Generation Sequencing techniques can be tricky for a non-programmer as it requires computational skills. iCOMIC serves as a robust platform for users with minimal programming skills. iCOMIC enables the user to choose from several pre-configured workflows for analyzing Whole Genome/Exome Sequencing and RNA-Seq data. Notably, iCOMIC also integrates novel algorithms developed in-house to predict cancer driver and passenger mutations, as well as tumor suppressor genes and oncogenes. The iCOMIC toolkit can be downloaded as a package and run on Linux, Windows, or Mac operating systems. iCOMIC enables easy analysis through an interactive GUI and hassle-free installation of software. The features listed above make iCOMIC an easily accessible, open-source pipeline for large genomic datasets. DNA-Seq benchmarking study performed in iCOMIC with GIAB dataset resulted in an F1 score of 0.971 and 0.988 for indels and SNPs, respectively. Similarly, for RNA Seq, comparison with a microarray dataset produced a fold change correlation coefficient of 0.85.

## Funding

This work was supported by the Department of Biotechnology, Government of India (DBT) (BT/PR16710/BID/7/680/2016), IIT Madras, Centre for Integrative Biology and Systems mEdicine (IBSE) and Robert Bosch Center for Data Science and Artificial Intelligence (RBCDSAI).

## Acknowledgements

The authors acknowledge Mr Likith Reddy for preliminary benchmarking studies and other members of the lab for useful discussions.

## Abbreviations

NGS: Next Generation Sequencing
GUI: Graphical User Interface
DNA-Seq: DNA Sequencing (Whole Genome/Exome Sequencing)
RNA-Seq: RNA Sequencing
iCOMIC: **i**ntegrating **CO**ntext of **M**utations **I**n **C**ancer.

## Notes

### Competing Interest Statement

The authors have declared no competing interest.

